# Multiple genetic pathways regulating lifespan extension are neuroprotective in a G2019S LRRK2 nematode model of Parkinson’s disease

**DOI:** 10.1101/2020.07.06.190025

**Authors:** Megan M. Senchuk, Jeremy M. Van Raamsdonk, Darren J. Moore

## Abstract

**Background:** Mutations in the *leucine-rich repeat kinase 2* (*LRRK2*) gene are the most frequent cause of late-onset, familial Parkinson’s disease (PD), and *LRRK2* variants are associated with increased risk for sporadic PD. While advanced age represents the strongest risk factor for disease development, it remains unclear how different age-related pathways interact to regulate *LRRK2*-driven late-onset PD.

**Findings:** In this study, we employ a *C*.*elegans* model expressing PD-linked G2019S LRRK2 to examine the interplay between age-related pathways and LRRK2-induced dopaminergic neurodegeneration. We find that multiple genetic pathways that regulate lifespan extension can provide robust neuroprotection against mutant LRRK2. However, the level of neuroprotection does not strictly correlate with the magnitude of lifespan extension, suggesting that lifespan can be experimentally dissociated from neuroprotection. Using tissue-specific RNAi, we demonstrate that lifespan-regulating pathways, including insulin/IGF-1 signaling, TOR, and mitochondrial respiration, can be directly manipulated in neurons to mediate neuroprotection. We extend this finding for AGE-1/PI3K, where pan-neuronal versus dopaminergic neuronal restoration of AGE-1 reveals both cell-autonomous and non-cell-autonomous neuroprotective mechanisms downstream of insulin signaling.

**Conclusions:** Our data demonstrate the importance of distinct lifespan-regulating pathways in the pathogenesis of *LRRK2*-linked PD, and suggest that extended longevity is broadly neuroprotective via the actions of these pathways at least in part within neurons. This study further highlights the complex interplay that occurs between cells and tissues during organismal aging and disease manifestation.

## Background

Parkinson’s disease (PD) is a common neurodegenerative disorder, affecting 1-2% of individuals over 65 years of age (de Lau and Breteler, 2006). With population demographics predicting an increasingly elderly population, the prevalence of PD is expected to grow to over 12 million cases by 2040 (GBD 2016 Neurology Collaborators, 2019; Dorsey et al, 2018). PD is a progressive movement disorder characterized by resting tremor, muscular rigidity, and bradykinesia primarily resulting from the selective degeneration of dopaminergic neurons in the substantia nigra and the corresponding loss of nigrostriatal pathway dopamine signaling. While PD is influenced by both genetic risk and environmental exposure, the etiology of PD remains unclear. Aging is considered a primary risk factor for the development of PD, as the incidence of disease rises exponentially with increasing age (Billingsley et al, 2018). Aging is associated with the impairment of numerous cellular pathways, including mitochondrial respiration, proteostasis, autophagy and stress responses that lead to a reduced chaperone response and increased genomic instability. A number of pathways have been identified that can regulate chronological age yet the relationship between these specific lifespan-regulating pathways and pathogenic mechanisms underlying PD are poorly understood.

The identification of genetic mutations linked to familial forms of PD have provided important insight into disease pathophysiology. Mutations in *leucine-rich repeat kinase 2* (*LRRK2*) cause late-onset, autosomal dominant familial PD that is largely indistinguishable from sporadic disease, whereas both coding and non-coding variants are associated with an increased risk of sporadic PD (Hernandez et al, 2016). *LRRK2* encodes a multidomain protein containing kinase, GTPase, and protein-protein interaction domains, with familial mutations commonly leading to enhanced kinase activity. The biological functions of LRRK2 remain to be fully elucidated, although prior studies suggest roles in diverse cellular processes, including vesicular transport/sorting, endocytosis, autophagy, mitochondrial dynamics, and lysosomal function. Familial LRRK2 mutations share the capacity to induce neuronal damage in culture models via a kinase-dependent mechanism, whereas increased LRRK2 kinase activity has been reported in PD-linked *D620N VPS35* knockin mice and sporadic PD brain. LRRK2 kinase activation may therefore represent a common mechanism underlying PD pathogenesis (Kluss et al, 2019), and a promising target for therapeutic disease-modification.

Among familial *LRRK2* mutations, G2019S is the most common, accounting for 5-6% of familial PD and 1-2% of sporadic PD in the US population (Billingsley et al, 2018). PD linked to the G2019S mutation is typically late-onset and exhibits variable yet age-dependent penetrance depending on ethnicity, with estimates of cumulative risk rising from ∼20% at 50 years to ∼75% at 80 years of age (Spatola and Wider, 2014). The exponential rise in PD risk after age 50 suggests that the aging process may play an important role in manifesting the pathogenic actions of *LRRK2* mutations (Marder et al, 2015). This interaction of age with LRRK2 is borne out in experimental models, such as G2019S LRRK2 knockin or transgenic mice, *Drosophila* or *C*.*elegans*, that develop age-dependent and often late-onset PD-like phenotypes (Longo et al, 2017; Ho et al, 2019; Xiong et al, 2017). Some individuals harboring *LRRK2* mutations escape PD within their lifetime, suggesting that genetic variation within different age-related pathways might feasibly be protective and could be manipulated to prevent disease.

While incomplete penetrance suggests that age-related pathways significantly interact with G2019S LRRK2 during disease progression, experimental mammalian models including human induced pluripotent stem cells and rodent models are cost-prohibitive or not practical for chronic aging studies. We therefore took advantage of the nematode *Caenorhabditis elegans*, a well-established model for aging studies, to systematically investigate functional interactions between age-related pathways and G2019S LRRK2-induced dopaminergic neurodegeneration. Conserved signaling pathways regulating the aging process include insulin/insulin-like growth factor 1 (IGF-1) signaling, target of rapamycin (TOR) pathway, signals from the reproductive system, environmental sensing, and caloric restriction. These pathways define major nutrient-sensing, endocrine, and stress response signaling pathways, and are required for the coordinated regulation of systemic organismal aging.

Our recent study demonstrated that reducing insulin/IGF-1 signaling, a well-characterized pathway for regulating lifespan, increased dopaminergic viability in *C*.*elegans* models of PD expressing human mutant LRRK2 or α-synuclein (Cooper et al, 2015). However, the underlying mechanism for this neuroprotective effect is not clear. It is possible that insulin/IGF-1 signaling interacts directly with G2019S LRRK2 to attenuate neurodegenerative phenotypes, and would therefore represent a potential therapeutic pathway. Alternatively, reduced insulin signaling may act indirectly by slowing the rate of aging in all tissues, and this feature may regulate LRRK2-induced neurotoxicity. To distinguish between lifespan extension in general versus the insulin signaling pathway as a key neuroprotective mechanism, we evaluate and compare the neuroprotective effects of multiple distinct lifespan-extending pathways in the G2019S LRRK2 worm model. Our data demonstrate that lifespan extension via multiple signaling pathways can regulate dopaminergic neuronal resilience towards G2019S LRRK2.

## Results

### G2019S LRRK2 induces age-dependent dopaminergic neuronal damage but does not influence lifespan

This study seeks to explore the role of different age-related pathways in regulating pathogenic phenotypes in a G2019S LRRK2 worm model of PD. We have employed a transgenic model expressing full-length human G2019S LRRK2 under the control of the presynaptic dopamine transporter promoter (Dat-1). A pDat-1::GFP transgene expressed in *cis* serves as a method to visualize the eight dopaminergic neurons in *C*.*elegans* from a total of ∼302 neurons. Previous studies have demonstrated that this model shows reduced dopaminergic neuronal function as indicated by subtle movement phenotypes and age-related changes in dopaminergic neurons monitored using pDat-1::GFP (Yao et al, 2010, Cooper al al, 2015). The pDat-1::hLRRK2(G2019S); pDat-1::GFP line (cwrIs856) is hereafter referred to as LRRK2(G2019S).

Our study focuses primarily on age-dependent neurodegenerative changes within the four anterior cephalic dopaminergic neurons (CEPs), as these neurons are relatively large in size with extended axonal processes and are simple to distinguish from intestinal autofluorescence (lipofusucin accumulation). Our initial validation of this model reveals that CEP neurons expressing LRRK2(G2019S) remain morphologically normal up to adult day 7 (Figure 1A). By adult day 9, large GFP-positive puncta are observed within and proximal to the CEP neuronal soma. By day 12, GFP-positive puncta are observed throughout the axon and dendrites, and GFP intensity within processes is reduced. Changes in cell soma morphology (i.e. swelling, reduced circularity) are also observed. As organismal aging continues to day 15, axonal processes become increasingly disorganized, and GFP intensity in the cell soma is greatly reduced. GFP appears to coalesce in puncta within the axon, as these remain bright despite the loss of GFP within the cell soma and dendritic processes. The GFP-positive puncta are consistent with the formation of axonal spheroids or inclusions that are commonly observed in degenerating neurons.

**Figure 1.**
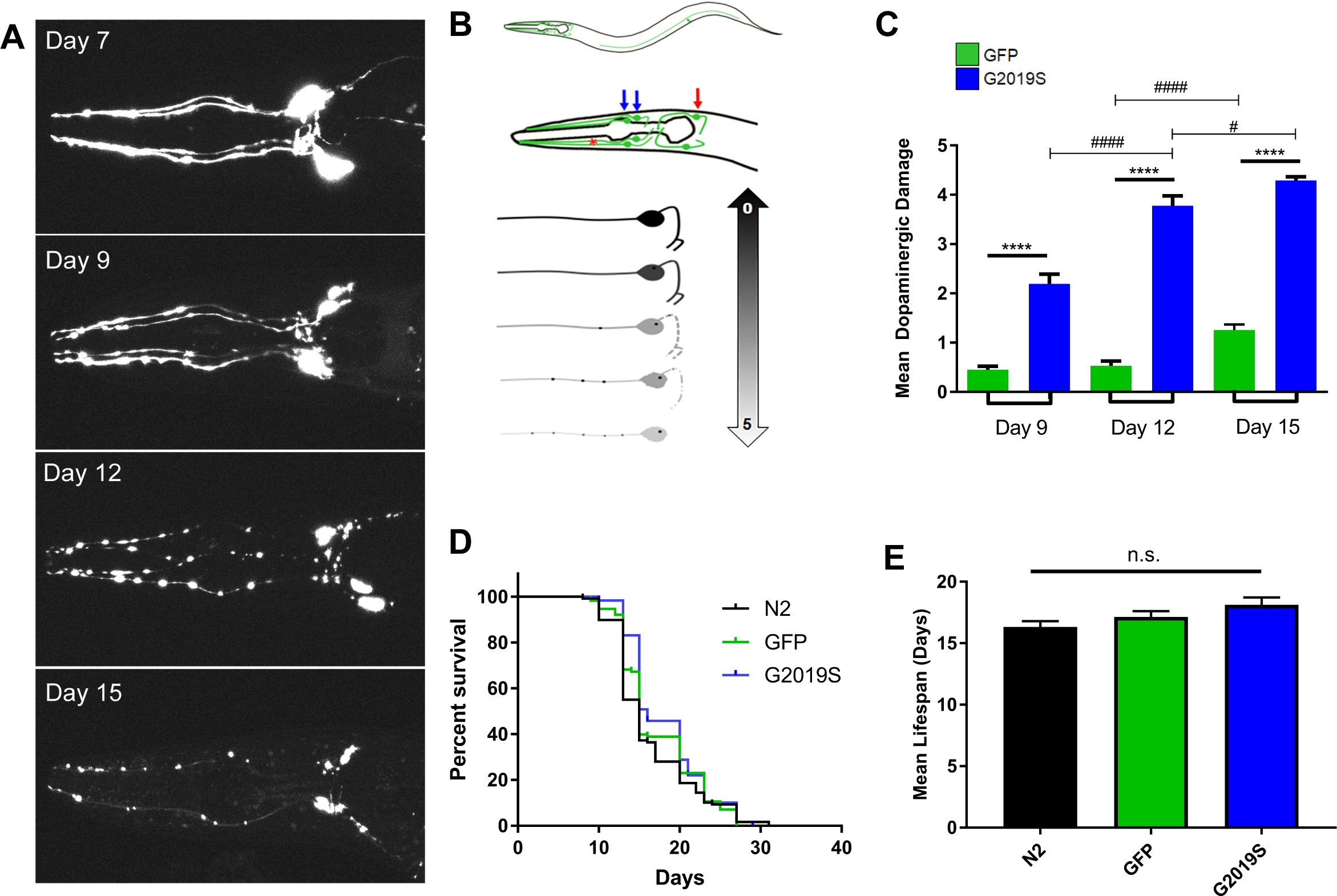
Human LRRK2(G2019S) expression in worms induces age-dependent dopaminergic neuronal damage without altering lifespan. **A**) Dopaminergic neurons are marked by GFP driven by the dopamine transporter (Dat-1) promoter. The pDat-1::hLRRK2(G2019S); pDat-1::GFP line (cwrIs856) is referred to as LRRK2(G2019S). Representative maximum projection confocal micrographs from LRRK2(G2019S) transgenic worms indicating increased GFP-positive dopaminergic neuronal damage during aging. Images are oriented with the mouth to the left. **B**) *C.elegans* dopaminergic signaling is carried out by 8 neurons from a total of 302 neurons. Two pairs of cephalic neurons (CEP, blue arrows) are the most anterior, followed by the anterior deirids (ADE, red arrows) located proximal to the nerve ring, and the laterally positioned posterior deirid pair (PDE). The axonal processes of the CEP neurons (red asterisk) extend toward the anterior, while the branched neuritic processes project distally into the synapse-rich nerve ring. The 0-5 scoring metric accounts for the overall morphology and damage of individual dopaminergic cell soma and neurites, the presence and localization of GFP-positive dense puncta (internal vs. external to cell body), and overall GFP intensity within individual neurons. **C**) Graph comparing dopaminergic neuronal damage to CEP neurons in worms expressing LRRK2(G2019S) or GFP alone at days 9, 12 and 15. Data represent dopaminergic damage with scoring from 0-5 (mean ± SEM; *n* ≥ 3 independent experiments). ^#^*P*<0.05 or ^####^*P*<0.0001 by one-way ANOVA with Tukey’s post-hoc test as indicated, or \*\*\*\**P*<0.0001 by unpaired Student’s *t*-test comparing GFP to LRRK2(G2019S) at each time point. LRRK2(G2019S) induces marked and progressive neuronal damage with age compared to GFP alone. **D-E**) LRRK2(G2019S) or GFP lines with transgene expression restricted to dopaminergic neurons (Pdat-1) exhibit a normal lifespan compared to wild-type N2 control worms as indicated by Kaplan-Meier survival analysis (**D**) and mean lifespan (**E**, mean ± SEM, *n* = 3 experiments, with 30 animals per experiment). **E**) One-way ANOVA finds no significant differences between mean lifespans of wild-type (N2, black), cwrIs730 [pDat-1::GFP] (green), and cwrIs856 [pDat-1::GFP; pDat-1::LRRK2(G2019S)] (blue) worm strains.

Based on these initial observations, we developed a robust scoring metric graded from 0 (normal) to 5 (maximum damage) to quantify overall damage in the four CEP dopaminergic neurons that incorporates measures of relative intensity of GFP, the formation of GFP-positive puncta in cell soma and processes, abnormal cell soma morphology (swelling, reduced circularity) and the disorganization of neuronal processes (Figure 1B). To further validate this scoring metric, we compare worms expressing pDat-1::GFP alone or together with LRRK2(G2019S) at days 9, 12 and 15 revealing that LRRK2(G2019S) expression induces pronounced dopaminergic neuronal damage at early time points with the accumulation of damage in an age-dependent manner compared to GFP alone (Figure 1C). A more modest reduction in GFP intensity and increased axonal disorganization are observed during aging in pDAT-1::GFP and pDAT-1::GFP; pDat-1::hLRRK2(WT) control lines compared to LRRK2(G2019S) (Figure 1C) (data not shown). However, GFP-positive puncta are never observed in these control lines, indicating that this phenotype is specific to LRRK2(G2019S). Despite dopaminergic neuronal damage induced by LRRK2(G2019S) expression, we observe that lifespan in this strain is not significantly altered compared to pDat-1::GFP or wild-type (non-transgenic N2) worms (Figure 1D-E), as observed previously (Yao et al, 2010; Cooper et al, 2015). Together, our data suggest that G2019S LRRK2 robustly induces age-dependent dopaminergic neuronal damage but without effects on lifespan in this worm model.

### Lifespan extension of LRRK2(G2019S) worms by genetic reduction of multiple negative regulators of longevity

To determine whether G2019S LRRK2-mediated neurodegeneration is impacted by alternative modes of lifespan extension, the LRRK2(G2019S) model was crossed to a well-characterized panel of genetic mutants known to extend lifespan (negative regulators of longevity, Table 1), including: *daf-2*, the insulin/IGF-1 receptor (Kenyon et al, 1993; Kimura et al, 1997); *age-1* (PI3K) (Friedman & Johnson, 1988), a downstream effector kinase in the insulin/IGF-1 signaling pathway; *nuo-6*, a mitochondrial complex I subunit (Yang & Hekimi, 2010); *glp-l* (NotchR), required for germline proliferation (Arantes-Oliveira et al, 2002); *eat-2*, a genetic model of caloric restriction (Lakowski & Hekimi, 1998); *rsks-1* (S6K), a downstream effector kinase of TOR signaling to regulate protein synthesis (Pan et al, 2007; Hansen et al, 2007); *ife-2* (eIF4E), a translation initiation factor (Hansen et al, 2007), and *osm-5*, an intraflagellar transport protein mediating chemosensation (Apfeld & Kenyon, 1999). Initially, we sought to confirm that lifespan would be extended in all mutant backgrounds. On a wild-type background, LRRK2(G2019S) transgenic worms exhibit a normal lifespan with an average of ∼17 days at 20°C similar to control wild-type (N2) worms (Figure 1D-E, Figure 2) (Yao et al, 2001; Cooper et al, 2015). When crossed into lifespan-extending mutant backgrounds, the mean lifespan of LRRK2(G2019S) worms is increased to a similar extent compared to each genetic mutant alone (Figure 2 and Table 1). Genetic mutants suppressing insulin/IGF-1 signaling (*daf-2* [e1370] and *age-1* [hx546]) have the strongest effect on lifespan extension, increasing mean lifespan to 39.1 days and 33.4 days of adulthood, respectively (Figure 2I). Lifespan is extended to a mean of 29.6 days in a *glp-1* mutant lacking germ cells, while mitochondrial respiration (*nuo-6*), TOR signaling (*rsks-1*), caloric intake (*eat-2*), and eIF4E-mediated protein translation (*ife-2*) mutants extend normal lifespan by ∼9 days over LRRK2(G2019S) alone or a wild-type N2 strain (Figure 2I). Reduction of chemosensory perception (*osm-5*) results in a more modest ∼20% increase in lifespan relative to control worms (Figure 2I). Overall, these data demonstrate that lifespan can be successfully extended using a diverse panel of genetic mutants in LRRK2(G2019S) transgenic worms similar to non-transgenic control worms.

**Table 1.** Comparison of lifespan extension induced by genetic mutants in wild-type and LRRK2(G2019S) worms

| Worm strain | Mean Lifespan (days) | Maximum Lifespan (days) | Mean Lifespan (% of N2 or G2019S) |
| --- | --- | --- | --- |
| Wild-type (N2) | 16.43 | 31 | - |
| **LRRK2(G2019S)** | **17.08** | **29** | - |
| *daf-2 (e1370)* | 39.28 | 60 | 239.07 |
| ***daf-2*; G2019S** | **39.06** | **57** | **228.69** |
| *age-1 (hx546)* | 30.81 | 49 | 187.52 |
| ***age-1*; G2019S** | **33.36** | **51** | **195.32** |
| *nuo-6 (qm200)* | 28.7 | 43 | 174.68 |
| ***nuo-6*; G2019S** | **26.56** | **44** | **155.50** |
| *rsks-1 (ok1255)* | 27.3 | 34 | 166.16 |
| ***rsks-1*; G2019S** | **26.01** | **34** | **152.28** |
| *eat-2 (ad1116)* | 24.65 | 36 | 150.03 |
| ***eat-2*; G2019S** | **26.43** | **38** | **154.74** |
| *glp-1 (e2141)* | 28.48 | 42 | 173.34 |
| ***glp-1*; G2019S** | **29.56** | **44** | **173.07** |
| *ife-2 (ok306)* | 23.55 | 35 | 143.34 |
| ***ife-2*; G2019S** | **24.63** | **40** | **144.20** |
| *osm-5 (ok451)* | 19.42 | 29 | 118.20 |
| ***osm-5*; G2019S** | **20.46** | **33** | **119.79** |

*For % mean lifespan, single genetic mutants were compared to N2 worms, whereas double mutants were compared to LRRK2(G2019S) (in bold).*

**Table 2.**
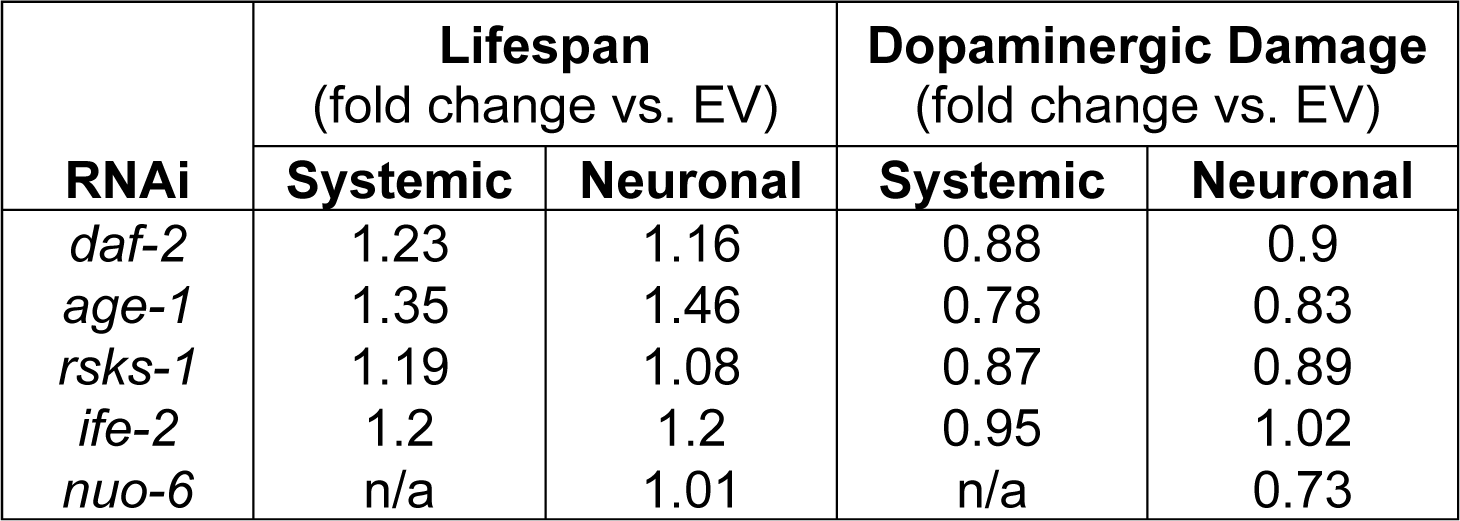
Comparison of fold change in lifespan and dopaminergic neuronal damage with systemic or neuronal-specific RNAi knockdown of longevity-regulating genes

**Figure 2.**
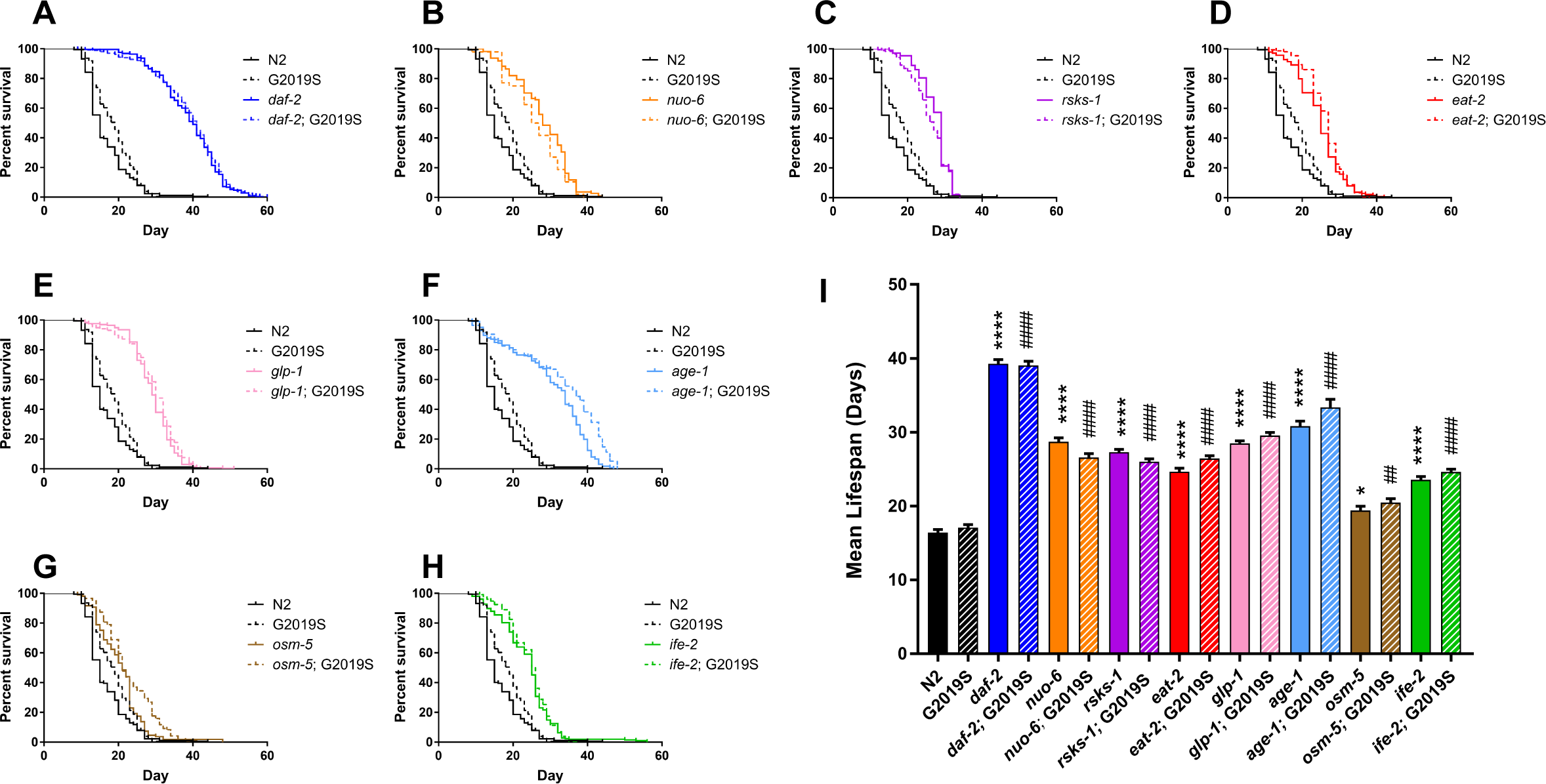
Genetic mutation of negative longevity regulators extends the lifespan of LRRK2(G2019S) transgenic worms. **A-H**) Kaplan-Meier survival curves comparing longevity of wild-type (N2, black solid) and LRRK2(G2019S) (black dashed) worms with lifespan-extending genetic mutants alone (color solid) and LRRK2(G2019S) transgenics on each mutant background (color dashed). Curves represent combined data from at least 3 individual experiments (*n* = 30 animals per experiment). **I**) Mean lifespan from data shown in **A-H**. Bars represent mean ± SEM. \**P*<0.05 or \*\*\*\**P*<0.0001 compared to WT (N2), ^##^*P*<0.01 or ^####^*P*<0.0001 compared to LRRK2(G2019S), by one-way ANOVA with Bonferroni’s multiple comparisons test.

### Lifespan-extending pathways provide neuroprotection against dopaminergic neuronal damage in LRRK2(G2019S) worms

We next assessed the extent to which different longevity pathways interact with LRRK2(G2019S)-induced neurotoxicity by monitoring damage to GFP-positive dopaminergic CEP neurons, comparing LRRK2(G2019S) transgenic worms on a wild-type background to lifespan-extending mutant backgrounds under identical conditions. Using the scoring metric validated in Figure 1, dopaminergic neuronal damage was assessed at day 14 of adulthood, a timepoint where degeneration is clearly observed but neurons largely remain intact. Intriguingly, we find that all pathways of lifespan extension confer variable neuroprotection in the LRRK2(G2019S) model with reduced scores of dopaminergic neuronal damage (Figure 3). In each of the eight lifespan-extending paradigms evaluated, CEP neuronal soma and axonal processes appear visibly healthier and brighter, with wider continuous processes that have increased GFP intensity and reduced GFP-positive puncta (Figure 3A). However, GFP-positive puncta are still detected in each mutant background, albeit at reduced levels, perhaps suggesting that mechanisms directly mediating LRRK2-induced toxicity are only partly impacted, and that overall organismal health is maintained for an extended period. Interestingly, we note that the degree of neuroprotection afforded by each mutant does not directly correlate with the degree of lifespan extension (Figure 2I and 3B). Linear regression analysis confirms the general trend that the extent of lifespan extension negatively correlates with degree of dopaminergic damage, as best exemplified by the opposing effects of *daf-2* and *osm-5* (Figure 3C). Certain lifespan-extending mutants, however, reveal that dopaminergic neuroprotection can be uncoupled from lifespan extension, resulting in an R^2^ value of 0.427. For example, *nuo-6* displays a moderate increase in lifespan yet shows pronounced reduction of dopaminergic damage, and likewise *age-1* shows robust lifespan extension yet a corresponding smaller degree of neuroprotection (Figure 3C). Collectively, these data demonstrate that delaying aging via multiple distinct pathways is broadly neuroprotective against G2019S LRRK2-induced neurotoxicity, yet lifespan extension does not strictly correlate with the magnitude of neuroprotection. Our data suggest that age-related pathways may functionally interact with G2019S LRRK2-dependent pathways to differing extents, specifically insulin signaling (*daf-2*), mitochondrial respiration (*nuo-6*) and TOR signaling (*rsks-1*), to mediate robust neuroprotection.

**Figure 3.**
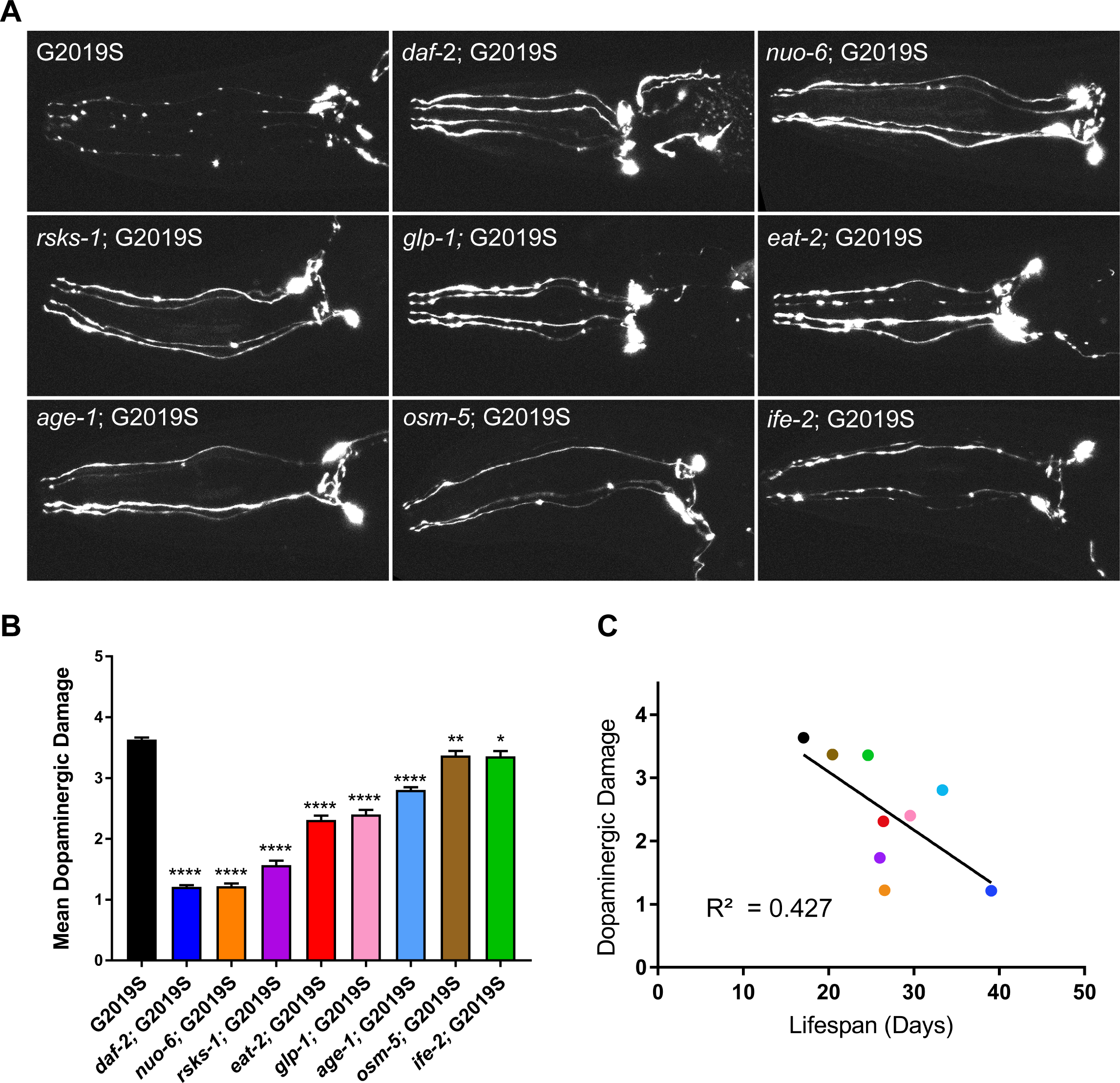
Lifespan extension is broadly neuroprotective in the LRRK2(G2019S) worm model. **A**) Representative maximum projection confocal images of anterior GFP-positive CEP dopaminergic neurons at day 14 of adulthood from LRRK2(G2019S) on a wild-type or genetic mutant background. Images reveal reduced yet variable neuronal damage in all lifespan-extending backgrounds. **B**) Quantification of CEP dopaminergic neuronal damage at day 14 using the 0-5 scoring metric. Bars represent the mean ± SEM damage from ≥3 individual experiments (*n* = 40 animals per experiment). \**P*<0.05, \*\**P<*0.005 or \*\*\*\**P*<0.0001 compared to LRRK2(G2019S) alone by one-way ANOVA with Dunnett’s post-hoc test. **C**) Linear regression analysis (R^2^ = 0.4266) of mean lifespan (in days) relative to mean dopaminergic neuronal damage (0-5 score) for each worm strain, reveals a general inverse correlation indicating that lifespan extension is broadly associated with reduced neuronal damage.

### Neuroprotective effects of *age-1* in the LRRK2(G2019S) model require DAF-16/FOXO

Neuronal health of LRRK2(G2019S)-expressing dopaminergic neurons is preserved in both the *daf-2* and *age-1* lifespan-extending backgrounds. AGE-1/PI3K functions as a downstream effector of the insulin receptor DAF-2 to control nuclear localization of the master transcriptional regulator DAF-16/FOXO. DAF-16/FOXO functions as a downstream factor required for a number of lifespan-regulating pathways, including nutrient and stress-related pathways (Insulin/IGF-1, TOR, caloric restriction, AMPK, JNK) as well as germline signaling. Accordingly, the ability of *daf-2* and *age-1* to increase lifespan are dependent on DAF-16 (Tissenbaum, 2018). We previously demonstrated that the protective effects of *daf-2* in LRRK2(G2019S) worms was dependent on the presence of DAF-16 (Cooper et al, 2015). We confirm that lifespan extension and dopaminergic neuroprotection in LRRK2(G2019S) worms conferred by *age-1*(*hx546*) similarly requires DAF-16 function by crossing *age-1*;LRRK2(G2019S) worms into a *daf-16* (*mu86*) mutant background (Figure 4). The introduction of *daf-16* effectively and fully reverses the increased lifespan (Figure 4A-B) and dopaminergic protective effects (Figure 4C) mediated by *age-1*. As the rate of aging is increased in worms with a *daf*-*16* mutation, dopaminergic damage was scored at day 10 as opposed to day 14. These data demonstrate that *age-1* and *daf-2* similarly require the canonical insulin/IGF-1 signaling pathway functioning upstream of DAF-16/FOXO for mediating neuroprotection in LRRK2(G2019S) transgenic worms.

**Figure 4.**
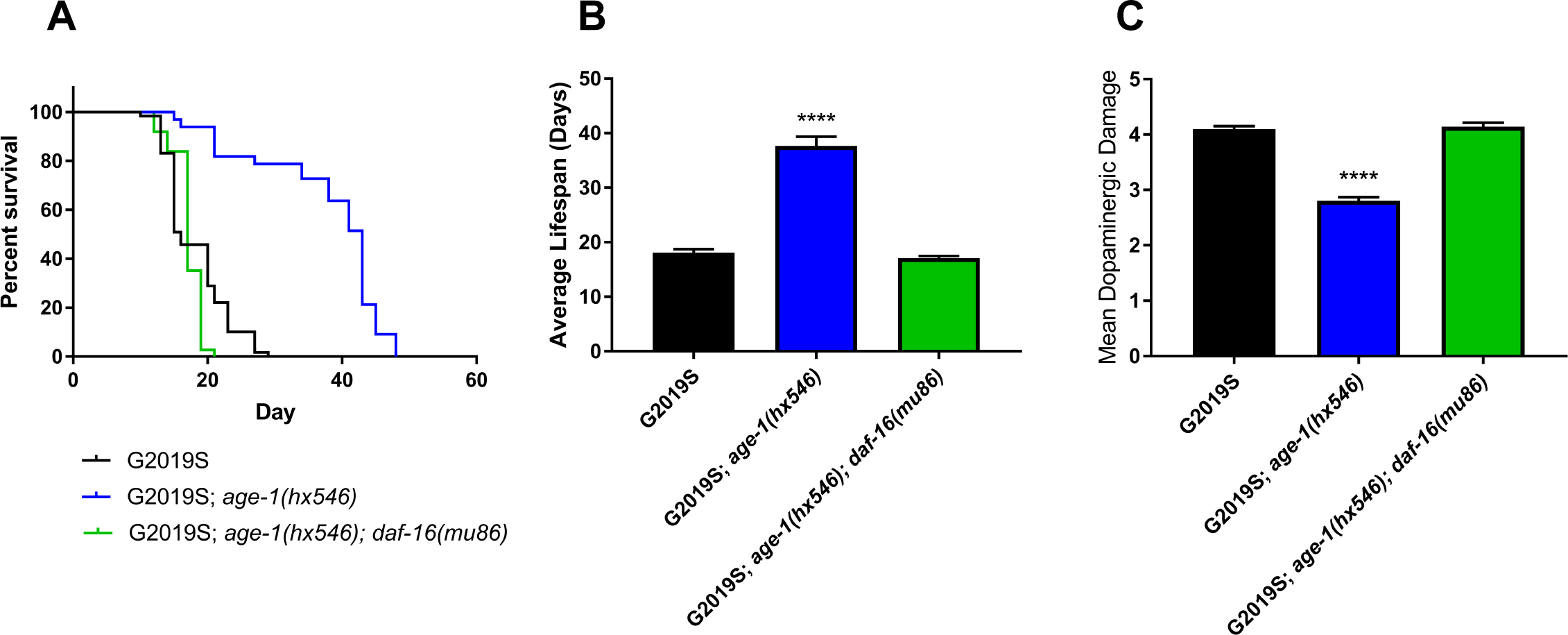
Lifespan extension and neuroprotection mediated by *age-1* mutant in LRRK2(G2019S) worms is dependent on DAF-16. Kaplan-Meier survival curve (**A**) and mean lifespan (**B**) of LRRK2(G2019S) worms on a wild-type background, and on an *age-1* or *age-1; daf-16* genetic background. The *daf-16* mutant fully reverses the extended lifespan conferred by *age-1*. **A**) Curves represent combined data from at least 3 individual experiments (*n* = 30 animals per experiment), whereas **B**) bars represent mean ± SEM. **C**) Quantitation of CEP dopaminergic neuronal damage at day 14 in LRRK2(G2019S) worms on all three genetic backgrounds. An *age-1* mutant markedly reduces neuronal damage compared to the wild-type background, that is fully restored to normal wild-type levels in an *age-1*; *daf-16* double mutant. Bars represent the mean ± SEM neuronal damage (*n* ≥ 3 independent experiments; with 40 animals/experiment). \*\*\*\**P*<0.0001 compared to LRRK2(G2019S) alone by one-way ANOVA with Dunnett’s post-hoc analysis.

### Lifespan-extending pathways mediate neuroprotection against G2019S LRRK2 in part via cell-autonomous effects in neurons

LRRK2(G2019S) is specifically expressed in dopaminergic neurons under the control of the Dat-1 promoter with damage limited to these neurons. The aging process requires the coordination of inputs from various tissues including the intestine, neurons and germline; input signaling can be perceived by one tissue but must be relayed to and acted upon by distal tissues. It is unclear whether lifespan-extending pathways mediate dopaminergic neuroprotection in LRRK2(G2019S) worms via their direct actions in neurons or indirectly via signals derived from peripheral tissues and organs. To address this question, we performed tissue-specific RNAi-mediated silencing of key lifespan-extending genes to compare the pan-neuronal versus body-wide effects of these pathways on lifespan-extension and neuroprotection against LRRK2(G2019S). We employed an RNAi targeting system using a dsRNA uptake channel SID-1 mutant (Systemic RNAi Deficient, *sid-1*[*qt9*]) combined with a rescuing SID-1 transgene driven by the pan-neuronal Unc-119 promoter (Calixto et al, 2010). In this system, neurons expressing the SID-1 transgene are selectively sensitized to RNAi versus non-neuronal cells that lack SID-1. Body-wide or systemic RNAi in a wild-type background has limited impact on gene expression in neurons as prior studies have suggested that *C*.*elegans* neurons are resistant to RNAi (Kamath et al, 2001; Timmons et al, 2001; Asikainen et al, 2005). The pDat-1::LRRK2(G2019S); pDat-1::GFP transgene was crossed into the neuronal-sensitizing pUnc-119::SID-1; *sid-1(qt9)* background, referred to hereafter as LRRK2(G2019S); SID-1. Relative to systemic body-wide RNAi in the LRRK2(G2019S) line, neuronal-specific RNAi in the LRRK2(G2019S); SID-1 background reveals a modest reduction in mean lifespan with control RNAi (empty vector) (Figure 5A-B).

**Figure 5.**
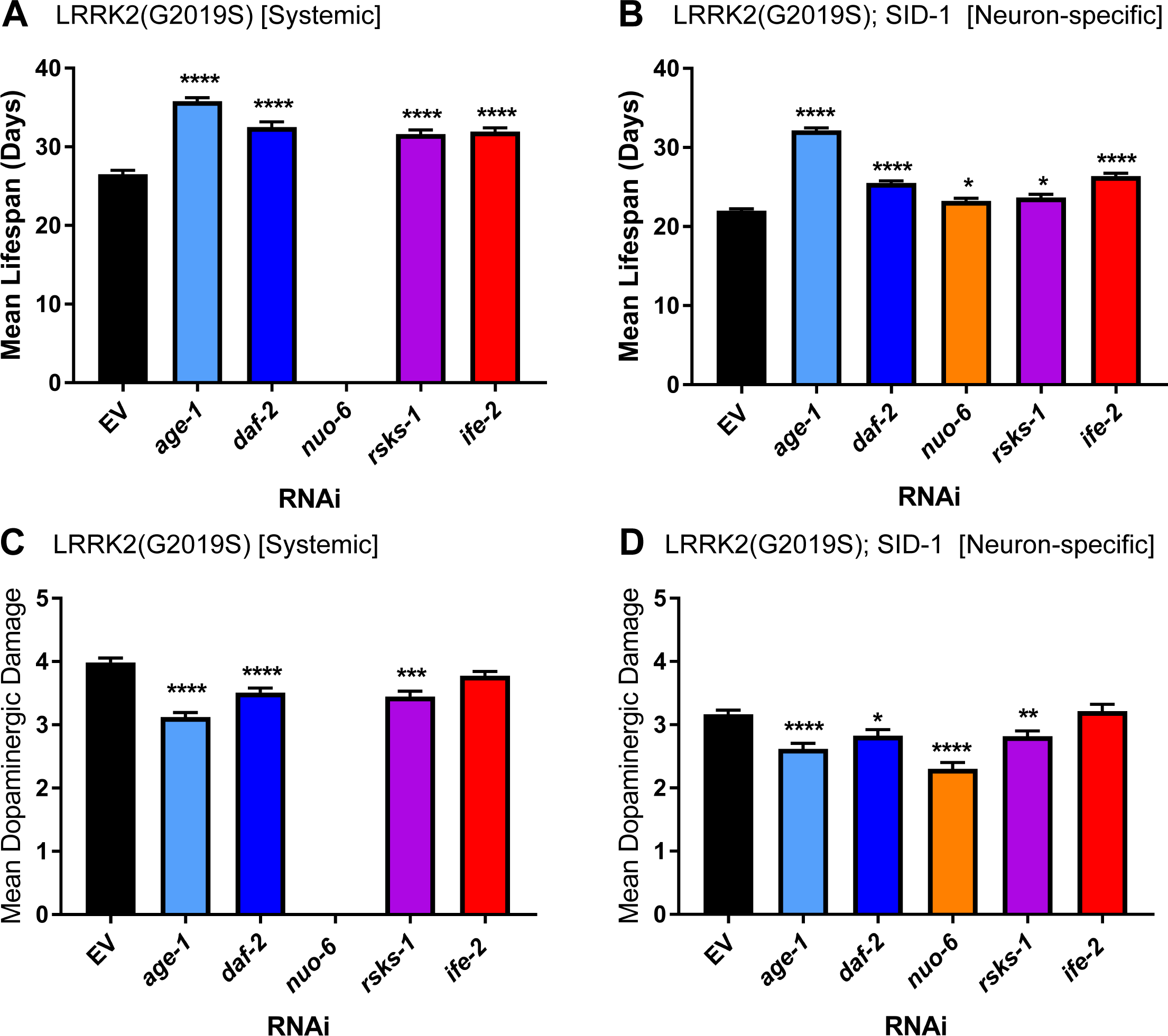
Neuronal-specific knockdown of multiple longevity-regulating genes is neuroprotective in the LRRK2(G2019S) worm model. A subset of negative regulators of longevity were targeted by RNAi-mediated knockdown either systemically (body-wide, with reduced efficacy in neurons) (**A, C**) or specifically within the neuronal cell population using the pUnc-119::SID-1; *sid-1*(pk3321) background (**B, D**). Mean lifespan of each strain is indicated (**A-B**). Quantitation of CEP dopaminergic neuronal damage in LRRK2(G2019S) worms with **C**) systemic body-wide RNAi assessed at day 14, or **D**) neuronal-specific RNAi assessed at day 11. Note: systemic knockdown of *nuo-6* results in early lethality and cannot be assessed in (**A, C**). Bars represent the mean ± SEM combined from ≥3 independent experiments with *n* ≥ 30 animals per experiment. \**P*<0.05, \*\**P*<0.01, \*\*\**P* <0.001 or \*\*\*\**P*<0.0001 compared to control (empty vector, EV) by one-way ANOVA with Dunnett’s post-hoc test.

Bacterial feeding clones from the Ahringer RNAi knockout library targeting lifespan-regulating pathways were verified by sequencing and first confirmed for lifespan extension phenotypes under systemic knockdown conditions prior to studies comparing systemic and neuron-specific RNAi. Clones for *eat-2* and *osm-5* were not available in the RNAi library. Instead, clones targeting related members of the chemoperception pathway (*che-3* and *osm-6*) were tested, but neither extended lifespan following RNAi-mediated knockdown and were not tested further (Figure S1). We find that the LRRK2(G2019S) [systemic RNAi] and LRRK2(G2019S); SID-1 [neuron-specific RNAi] backgrounds both exhibit marked lifespan extension in response to RNAi targeting the insulin/IGF-1 pathway (*daf-2, age-1*) (Figures 5A-B and S2). Surprisingly, neuron-specific RNAi to *age-1* results in a 1.46-fold increase in lifespan similar to systemic RNAi (1.35-fold), suggesting that the lifespan-regulating functions of AGE-1/PI3K are mediated primarily via the neuronal population. Interestingly, RNAi for *daf-2* demonstrates a stronger increase in lifespan with systemic (1.23-fold) compared to neuronal (1.16-fold) knockdown. These data are in agreement with previous studies demonstrating that pan-neuronal restoration of insulin/IGF-1 function partially rescues extended lifespan in mutant backgrounds (Wolkow et al, 2000; Iser et al, 2007). Importantly, the dopaminergic degenerative phenotype in LRRK2(G2019S) worms is suppressed by systemic or neuronal RNAi for *age-1* and *daf-2* (Figure 5C-D), similar to data obtained with genetic mutants (Figure 3), thereby suggesting that the insulin/IGF-1 signaling pathway can interact with aging and neurodegeneration via both cell autonomous and non-cell autonomous functions.

Similar to the insulin/IGF-1 pathway, targeting *rsks-1* by systemic or neuronal-specific RNAi induces lifespan extension (Figure 5A-B and S2), suggesting longevity-regulating functions can occur in both neuronal and non-neuronal tissues. The pan-neuronal or systemic reduction of *rsks-1* function reduces dopaminergic neuronal damage in LRRK2(G2019S) worms to a similar extent (Figure 5C-D), although TOR pathway signaling has a greater impact on lifespan in peripheral tissues (1.2-fold) versus neurons (1.1-fold). Systemic RNAi for *nuo-6* results in early larval arrest and embryonic lethality (Figure 5A and S2), as observed previously (Yang et al, 2010; Sonnichsen et al, 2005; Rual et al, 2004; Fraser et al, 2000), that precludes further analysis. However, neuronal-specific RNAi for *nuo-6* results in modest lifespan extension (Figure 5B and S2) yet a marked reduction in dopaminergic neuronal damage (Figure 5D) in LRRK2(G2019S) worms. These data suggest a direct role for mitochondrial respiration in LRRK2(G2019S)-induced dopaminergic phenotypes that can be uncoupled from mechanisms associated with lifespan extension induced by mitochondrial dysfunction. In contrast, systemic or neuronal-specific knockdown of *ife-2* induces marked lifespan extension (Figure 5A-B and S2) but fails to protect against G2019S LRRK2-induced dopaminergic damage (Figure 5C-D), demonstrating that neuroprotection does not strictly correlate with extended longevity. Collectively, our data suggest that age-related pathways can directly function in part within neurons (*age-1, daf-2, nuo-6, rsks-1*) to mediate lifespan extension and dopaminergic neuroprotection in LRRK2(G2019S) worms.

### Restoration of AGE-1/PI3K in neurons differentially modulates lifespan and dopaminergic neuronal damage in LRRK2(G2019S); *age-1* worms

Using tissue-specific RNAi, we demonstrate that a number of lifespan-regulating pathways mediate systemic effects on longevity via signaling occurring within neuronal populations, as well as exerting lifespan-regulating effects in peripheral non-neuronal tissues and cells. It is possible that reduced LRRK2(G2019S)-induced dopaminergic damage observed in lifespan-extending conditions can be attributed to systemic pathways regulating organismal aging, as both insulin/IGF-1 signaling and mitochondrial dysfunction can function via non-cell-autonomous mechanisms (Wolkow et al, 2000; Iser et al 2007; Libina et al, 2003; Zhang et al, 2013; Apfeld & Kenyon 1998). Alternatively, signaling could be required within specific cells or populations to directly regulate their viability and health. To address whether insulin/IGF-1 signaling functions specifically within dopaminergic neurons to impact LRRK2(G2019S)-induced neurotoxicity, we compared the extent of neuroprotection in LRRK2(G2019S); *age-1* worms with restoration of AGE-1 specifically targeted to dopaminergic neurons or pan-neuronally. We generated extrachromosomal transgenic arrays that drive expression of AGE-1/PI3K cDNA under the control of the pan-neuronal Unc-119 promoter or the dopaminergic-specific Dat-1 promoter, that were crossed into long-lived LRRK2(G2019S); *age-1*(hx546) worms for rescue studies. We elected to focus on the AGE-1/PI3K pathway for these studies since RNAi targeting of *age-1* in neurons produces robust effects on both lifespan and neuroprotection compared to other pathways suggesting a prominent role in neurons (Figure 5). In these new lines, lifespan and dopaminergic neuronal damage were monitored.

Pan-neuronal (pUnc-119-driven) expression of AGE-1 is able to rescue lifespan extension conferred by *age-1*(hx546), while dopaminergic-specific AGE-1 restoration does not impact longevity (Figure 6A-B). These data demonstrate that AGE-1 function is required in neurons in general to regulate lifespan, as previously suggested (Iser et al, 2007; Wolokow et al, 2000), whereas AGE-1 in dopaminergic neurons does not participate in lifespan. At day 14, dopaminergic neuronal damage in LRRK2(G2019S) worms is pronounced (score = 4.1 ± 0.06), and the introduction of *age-1* rescues neuronal damage (score = 2.81 ± 0.06) (Figure 6C-D). Restoration of AGE-1 expression in dopaminergic neurons (pDat-1::AGE-1) is sufficient to significantly and partially restore neuronal damage induced by LRRK2(G2019S) (score = 3.28 ± 0.07) (Figure 6C-D). These data suggest that interactions between AGE-1 and LRRK2(G2019S) can occur cell-autonomously within dopaminergic neurons, despite also having a non-cell-autonomous role in lifespan regulation. Surprisingly, pan-neuronal restoration of AGE-1 further enhances protection (reduces damage) in dopaminergic neurons (score = 2.34 ± 0.08) beyond the protective effects of *age-1* alone (Figure 6C-D). Similar results with respect to lifespan extension and dopaminergic neuronal damage are observed with at least two independent array lines expressing AGE-1 (Figure S3). These complex data suggest that dopaminergic neuronal health may be regulated directly by cell-autonomous AGE-1 signaling, as well as indirectly by non-cell-autonomous inputs arising from outside the nervous system. The observation that expressing AGE-1 in neurons of LRRK2(G2019S); *age-1* worms decreases lifespan but increases neuronal survival demonstrates that lifespan and neuroprotection can be experimentally dissociated.

**Figure 6.**
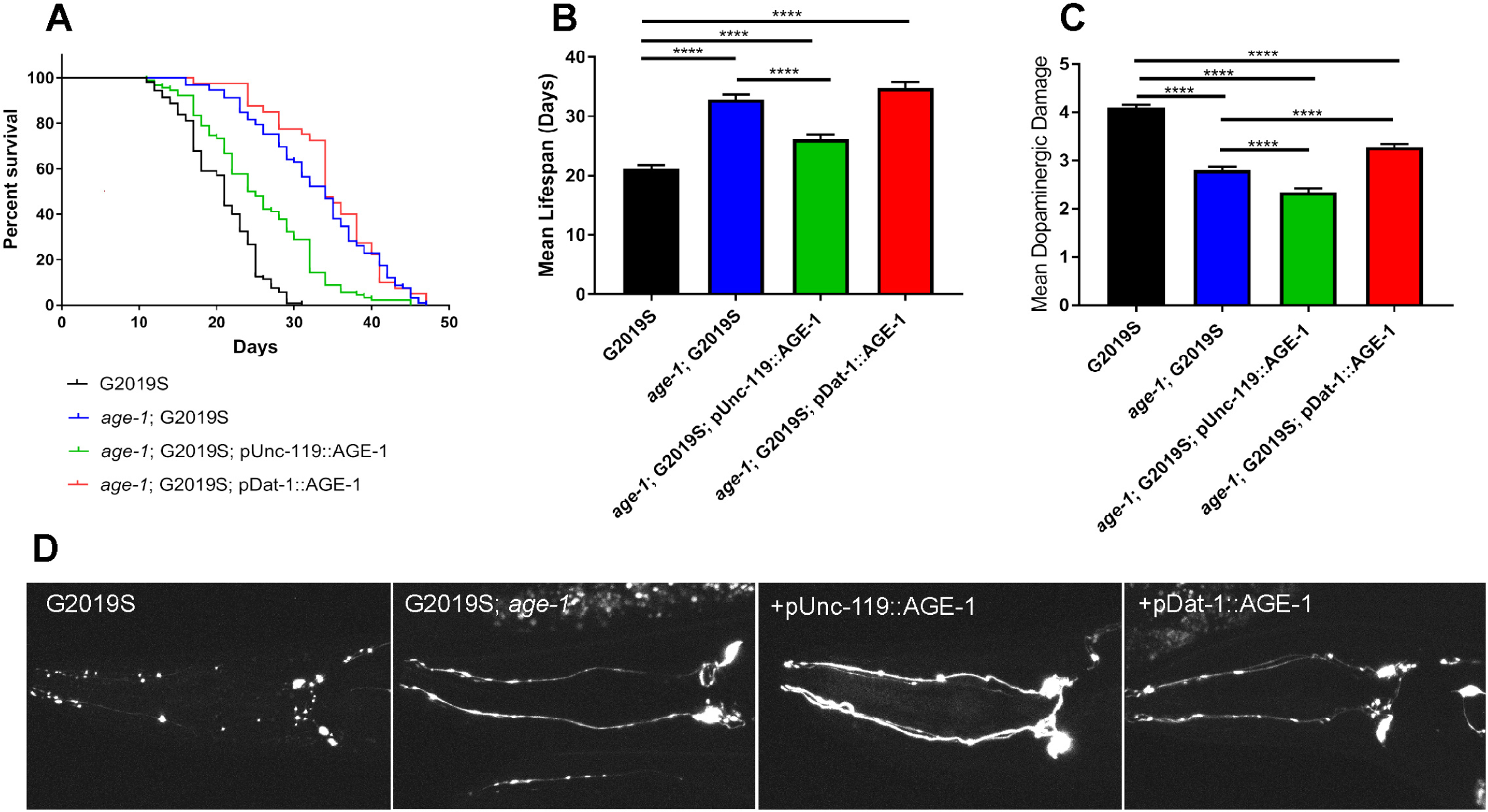
Restoring AGE-1/PI3K function in dopaminergic neurons demonstrates a cell-autonomous effect on LRRK2(G2019S)-induced neuronal damage. **A**) Kaplan-Meier survival curves comparing the lifespan of LRRK2(G2019S) (black), *age-1*; LRRK2(G2019S) (blue), and transgenic lines expressing AGE-1 from pan-neuronal (pUnc-119; green) or dopaminergic-specific (pDat-1; red) promoters in the *age-1*; LRRK2(G2019S) background. **B**) Mean lifespan or each worm strain from survival data shown (**A**). Extended lifespan in the *age-1* background is partially reverted by pan-neuronal AGE-1 expression. **C**) Quantification of mean dopaminergic neuronal damage induced by LRRK2(G2019S) at day 14 in wild-type, *age-1* mutant, or in tissue-specific AGE-1 rescue lines on an *age-1* mutant background. **B-C**) Bars represent the mean ± SEM from ≥3 independent experiments with *n* ≥ 30 animals per experiment. \*\*\*\**P*<0.0001 between the indicated groups by one-way ANOVA with Dunnett’s multiple comparisons post-hoc test. **D**) Representative maximum projection confocal images of GFP-positive CEP dopaminergic neurons at day 14 in pDat-1::GFP; pDat-1::LRRK2(G2019S) worms on wild-type, *age-1* or transgenic AGE-1 rescue backgrounds.

## Discussion

Understanding LRRK2 pathophysiological mechanisms is critical for defining cellular pathways and targets that are amenable to therapeutic intervention for attenuating neurodegeneration in *LRRK2*-linked PD. In this study, we evaluate how distinct mechanisms regulating the aging process interact with human LRRK2(G2019S)-induced dopaminergic neurodegeneration, addressing whether longevity is intrinsically coupled to dopaminergic neuronal health. We first demonstrate that mean lifespan of LRRK2(G2019S) worms can be successfully extended by different age-related pathways, including reduction of insulin/IGF-1 signaling (*daf-2*; *age-1*), TOR signaling (*rsks-1*), caloric intake (*eat-2*), chemoperception (*osm-5*), germline function (*glp-1*) or mitochondrial function (*nuo-6*). Next, we compared how LRRK2(G2019S)-induced dopaminergic degeneration was impacted by genetic mutants with different levels of lifespan extension. We find that age-dependent neuronal damage induced by LRRK2(G2019S) is broadly regulated by distinct lifespan-extending pathways, with extended longevity generally correlating with dopaminergic neuroprotection. Importantly, our data demonstrate that neuroprotection against LRRK2(G2019S) is not simply limited or specific to *daf-2*, as we previously reported (Cooper et al, 2015), but extends beyond insulin-IGF-1 signaling. However, we find that lifespan-extending paradigms variably influence the extent of neuroprotection with a non-linear relationship between lifespan and dopaminergic damage in the LRRK2(G2019S) model. These data suggest that lifespan extension *per se* is not sufficient alone for protection, and supports instead a specific interaction between certain age-related pathways and LRRK2(G2019S)-related pathogenic mechanisms, including most prominently insulin/IGF-1 signaling, TOR-mediated protein translation, and mitochondrial respiration. Our study highlights these age-related pathways as important for *LRRK2*-linked disease that warrant further mechanistic studies in mammalian LRRK2 models.

The tissue specificity mediating the interaction between LRRK2(G2019S) and lifespan-regulating factors was examined using neuronal-specific versus body-wide (non-neuronal) RNAi-mediated gene silencing. Neuronal-specific or body-wide knockdown of *age-1, daf-2, rsks-1*, and *nuo-6* are sufficient to induce lifespan extension and reduce LRRK2(G2019S)-induced dopaminergic damage, suggesting roles for these pathways in both neuronal and peripheral non-neuronal cells. However, one clear example of lifespan extension being experimentally dissociated from neuroprotection is provided by *ife-2* knockdown that extends lifespan with body-wide or neuronal-specific RNAi yet fails to impact dopaminergic degeneration. Interestingly, we find that knockdown of *nuo-6* that induces lifespan extension via mitochondrial dysfunction (Yang et al, 2010), interacts with LRRK2(G2019S), as both a systemic genetic mutant and neuronal-specific RNAi lead to a modest increase in lifespan yet a dramatic reduction of LRRK2(G2019S)-induced neurotoxicity. *Nuo-6* therefore provides another example of this dissociation that may imply this mitochondrial-related pathway rather than lifespan extension most likely drives the protective interaction with LRRK2(G2019S). The loss of *nuo-6* function, and impaired mitochondrial complex-I activity, appears to contradict long-held observations that mitochondrial impairment contributes to PD (Singh et al, 2019). It is important to note that *nuo-6* genetic mutation or body-wide RNAi both induce early lethality in worms, but not when limited to neurons, implying that the beneficial effects of lowering *nuo-6* arise solely from neurons. Mitochondrial dysfunction is sensed and adapted to early in *C*.*elegans* development (Dillin et al, 2002), leading to upregulation of stress response and other related pathways that may specifically protect against LRRK2(G2019S) later in life. It is most likely therefore that these stress-related adaptations in neurons rather than complex-I deficits induced by reduced *nuo-6* levels are sufficient to mediate neuroprotection.

Our study also focused on the insulin/IGF-1 signaling pathway as a mechanism of delayed aging and increased neuroprotection in the LRRK2(G2019S) model of PD. Reducing insulin receptor signaling or increasing DAF-16/FOXO activity specifically in adipose or neuroendocrine tissue has been shown to increase lifespan in worms, flies and mice (Apfeld and Fontana, 2018), and has been shown to have both cell-autonomous and non-cell-autonomous roles in the regulation of lifespan and other metabolic processes. In agreement with previous studies (Tank et al, 2011; Toth et al, 2010; Pan et al, 2011; Wolkow et al, 2000; Iser et al, 2007), we find that the insulin/IGF-1 signaling pathway is involved in nervous system aging, including preservation of dopaminergic neuronal integrity and survival. We find that specifically restoring AGE-1/PI3K function in dopaminergic neurons is sufficient to partially reverse the neuroprotective effects against LRRK2(G2019S) mediated by *age-1* disruption. However, pan-neuronal restoration of AGE-1 activity, which is capable of reversing lifespan extension in systemic *age-1* mutants, appears to have an opposite effect on dopaminergic neurodegeneration induced by LRRK2(G2019S). This unexpected finding demonstrates that the insulin/IGF-1 signaling pathway likely acts in both a cell-autonomous (i.e. dopaminergic) and non-cell-autonomous manner with respect to neuronal health, perhaps through the activation of stress response pathways in non-dopaminergic neurons and/or peripheral tissues that elicit further neuroprotection against LRRK2(G2019S). This observation highlights the importance of studying neurodegeneration and potential therapeutic interventions in the context of the whole organism.

## Conclusions

Our study provides insight into the pathogenesis of *LRRK2*-linked PD and longevity regulation. We find that LRRK2(G2019S)-induced neurodegeneration is strongly influenced by distinct pathways governing the overall rate of aging and lifespan, including insulin/IGF-1 signaling, the TOR pathway, and mitochondrial respiration. These and other lifespan-regulating pathways are highly conserved and associate with major signaling pathways. While distinct sets of factors are engaged downstream, mechanisms of lifespan modulation at least partially overlap, and cross-talk between signaling pathways may partly account for the observed complexity. LRRK2 has been shown to interact with different cell signaling pathways, specifically those involved in protein degradation, mitochondrial and endolysosomal functions. Further examination of the interplay between these pathways may reveal common factors influencing LRRK2-dependent neurotoxicity, and future studies will require validation of these mechanisms in genetic rodent models of PD. Overall, our study highlights that systemic aging of the organism and neuronal degeneration are related phenomena, but in a non-linear manner, and suggests that insulin/IGF-1 signaling is compartmentalized within different cells and tissues of an organism that are potentially differentially regulated during aging. Taken together, our study highlights specific lifespan-regulating pathways that may provide a number of promising therapeutic targets for inhibiting LRRK2-dependent neurodegeneration in PD.

## Methods

### *C.elegans* strains and maintenance

Unless noted, strains were maintained at 20°C on NGM seeded with OP50 bacteria according to standard practice (Brenner, 1974). Transgenic cwrIs856 [Pdat-1::GFP, Pdat-1::LRRK2(G2019S), lin-15(+)] has been previously described (Yao et al, 2010; Cooper et al, 2015). Double mutants were constructed as detailed elsewhere (Cooper et al, 2015). *daf-2* animals were raised at 15°C and shifted to 20°C at L4 stage. *glp-1* animals were raised at 15°C and shifted to 25°C as embryos to obtain germline(−/−) adults. cwrIs856 [pDat-1::GFP; pDat-1::LRRK2(G2019S)] were treated similarly in all conditions to serve as controls. For all assays, a minimum of three biological replicates (∼40 worms) per strain were assayed. All assays were performed by an experimenter blinded to genotype.

### Lifespan analysis

Worm populations were synchronized by washing a well-seeded plate to remove adults and larvae, leaving a population of embryos. After 2-3 days, upon reaching young adulthood, 30-40 worms were transferred to NGM plates containing 25 µM fluorodeoxyuridine (FUdR); inclusion of this DNA synthesis inhibitor prevents hatching of progeny following the initial transfer (at 48 h post-plating). FUdR has minimal effects on longevity at this concentration (Van Raamsdonk & Hekimi, 2011), and does not affect expression levels or localization of GFP transgenes.

Animals were transferred to fresh plates weekly, and viability was scored every 2 days by gentle prodding with a platinum pick. Animals that failed to respond were scored as dead. Worms that died from internal hatching of progeny, expulsion of internal contents or desiccation on the side of the dish were censored and excluded from analysis.

### Dopaminergic neuronal damage

Dopaminergic neurons were marked by GFP expression driven by the dopamine transporter (DAT-1) promoter. To quantify damage and compare across treatments, we developed a scoring metric of 0-5, with a score of 0 given to healthy, unbroken, bright, thick axons and cell bodies lacking punctate structures, as these are not observed in pDat-1::GFP-expressing strains that lack the LRRK2(G2019S) transgene. A score of 5 was given to neurons severely damaged (multiple puncta and/or cell body morphology defects) or undetectable GFP. Transmission electron microscopy has been used previously to demonstrate that loss of GFP in these neurons correlates with ultrastructural damage consistent with apoptotic cell death by (Nass et al, 2002).

Quantitative analyses of dopaminergic degeneration was performed by immobilizing 20-40 worms (unless stated otherwise, aged to Day 14 on 25 µM FUdR) on a 1.5% agarose pad with 5 mM levamisole. Live worms on coverslipped slides were immediately scored at 40x using a Leica upright epifluorescent microscope (DM5500 B). Unless noted otherwise, all images are maximum projection composite micrographs collected on a Nikon A1plus-RSi scanning confocal microscope. Images were collected every 0.5 µm over 30 µm, capturing the cell body and axonal projections of pDat-1::GFP-labeled cephalic sensilla (CEP) neurons. All images shown are representative of multiple independent experiments.

### RNAi

Standard NGM agar was supplemented with 50 µg/ml carbenicillin and 5 mM isopropyl β-D-1-thiogalactopyranoside (IPTG) post-autoclaving and allowed to dry overnight at room temperature. RNAi cultures were prepared by sequence verifying single colony cultures from the Ahringer RNAi plasmid library. *E.coli* HT115 carrying sequence-verified RNAi targeting plasmid clones were grown in LB supplemented with 50 µg/ml carbenicillin and grown for 8 hours at 37°C. Cultures were concentrated 5x prior to seeding plates with 200 µl of RNAi bacteria, which was spread, dried, and allowed to induce dsRNA expression for 48 hours at RT, after which point plates were stored at 4°C for up to 1 week. HT115 carrying the pL4440 plasmid lacking a gene-targeting cassette was used as a control for all RNAi experiments (empty vector).

RNAi knockdown was initiated by plating L4 P_o_ animals to prepared plates. After 30-35 h, adult animals were transferred to fresh RNAi plates and allowed to lay for 12 h, after which point adults were removed. After an additional ∼50 h, when embryos laid on RNAi plates had developed to young adulthood, animals were transferred to fresh RNAi plates containing 25 µM (FUdR) in addition to carbenicillin and IPTG.

Neuron-specific RNAi was performed by crossing TU3401 (*sid-1*(pk3321); uIs69[pCFJ90(myo-2p::mCherry) + unc-119p::sid-1]) (Calixto et al, 2010) to cwrIs856 [pDat-1::GFP; pDat-1::LRRK2(G2019S)]. Lifespan and neuronal damage were assayed as above, however, due to the modestly shortened lifespan attributed to *sid-1*(pk3321) in the periphery, dopaminergic neuronal damage was assayed at Day 11, a point at which control animals show equivalent damage to non-*sid-1* animals at Day 14.

### Construction of AGE-1 transgenics

Tissue-specific rescue transgenic expression constructs of the AGE-1 ORF were generated by standard Multi-site Gateway cloning methods. Full-length *age-1* cDNA was PCR-amplified using Phusion High-Fidelity DNA Polymerase+GC buffer (Thermofisher) using total cDNA prepared from N2 mRNA (Ampliscribe). AttB1 and AttB2 adaptors were added to the 5’ end of the forward and reverse primers, respectively, to allow for BP-mediated recombination into a pDONR221 vector (Invitrogen). The resulting entry clone was sequence verified. LR recombination (Invitrogen) was used to generate AGE-1 flanked by a 5’ promoter element (either pan-neuronal pUnc-119 or dopaminergic-specific pDat-1, both from Dharmacon), and a 3’UTR (pADA-126; let-858 3’UTR in pDONRP2R-P3) in the pCFJ150 MosSCI destination vector. Resulting expression plasmids were confirmed by diagnostic digest. Injection mixes were prepared by digesting pMS16 (pUnc-119::AGE-1::let-858) and pMS17 (pDat-1::AGE-1::let-858) with AgeI and Srf1; the resulting ∼11.6 kb band was gel purified and injected into N2 animals at 5 ng/µl with co-marker (pMyo-3::RFP (bsem1122) at 5 ng) and carrier DNA (sheared salmon testis DNA) to 100 ng/µl. The presence of the AGE-1 ORF transgene was verified in RFP positive lines by PCR. AGE-1/RFP double-positive arrays were then crossed to *age-1*(hx546); cwrIs856 [pDat-1::GFP; pDat-1::LRRK2(G2019S)] to generate tissue-specific restoration lines. Homozygosity of the *hx546* missense mutant was confirmed using intron-specific primers, allowing for genomic locus-specific sequencing.

### Statistical analysis

All statistical significance was determined using GraphPad Prism. Lifespan assays were assessed using the log-rank test to compare survival curves. Statistical significance of differences of lifespan means and dopaminergic neuronal damage was determined using a one-way ANOVA with Bonferroni or Tukey post-hoc tests to identify specific differences. Data are presented as mean ± standard error of the mean (SEM).

### *C.elegans* strains used in this study

*C.elegans* strains used in this study:
| Strain | Genotype | Original publication |
| --- | --- | --- |
| SGC730 | cwrls730 [Pdat-1::GFP, lin-15(+)] | Yao et al, 2010 |
| SGC722 | cwrls722 [Pdat-1::GFP, Pdat-1::LRRK2(WT), lin-15(+)] | Yao et al, 2010; Cooper et al, 2015 |
| CB1370 | daf-2(e1370) III | Kenyon et al, 1993; Kimura et al, 1997 |
| TJ1052 | age-1(hx546) II | Friedman & Johnson, 1988 |
| MQ1333 | nuo-6(qm200) I | Yang & Hekimi, 2010 |
| CB4037 | glp-1(e2141) III | Priess et al., 1987;<br>Arantes-Oliveira et al., 2002 |
| DA1116 | eat-2(ad1116) II | Lakowski & Hekimi, 1998 |
| KX15 | ife-2(ok306) X | Syntichaki et al, 2007 |
| VC265 | osm-5(ok451) X | Apfeld and Kenyon, 1999 |
| RB1206 | rsks-1(ok1255) III | Hansen et al, 2006; Pan et al, 2007 |
| JVR202 | daf-2(e1370) III; cwrls856 [Pdat-1::GFP, Pdat-1::LRRK2(G2019S), lin-15(+)] | Cooper et al, 2015 |
| DJM050 | age-1(hx546) II; cwrls856 [Pdat-1::GFP, Pdat-1::LRRK2(G2019S), lin-15(+)] |  |
| JVR321 | nuo-6(qm200); cwrls856 [Pdat-1::GFP, Pdat-1::LRRK2(G2019S), lin-15(+)] |  |
| DJM049 | glp-1(e2141); cwrls856 [Pdat-1::GFP, Pdat-1::LRRK2(G2019S), lin-15(+)] |  |
| JVR320 | eat-2(ad1116) II.; cwrls856 [Pdat-1::GFP, Pdat-1::LRRK2(G2019S), lin-15(+)] |  |
| JVR315 | ife-2(ok306) X.; cwrls856 [Pdat-1::GFP, Pdat-1::LRRK2(G2019S), lin-15(+)] |  |
| DJM048 | osm-5(ok451); cwrls856 [Pdat-1::GFP, Pdat-1::LRRK2(G2019S), lin-15(+)] |  |
| DJM068 | rsks-1(ok1255) III; cwrls856 [Pdat-1::GFP, Pdat-1::LRRK2(G2019S), lin-15(+)] |  |
| DJM067 | daf-16(mu86) I; age-1(hx546) II; cwrls856 [Pdat-1::GFP, Pdat-1::LRRK2(G2019S), lin-15(+)] |  |
| DJM063 | age-1(hx546) II; cwrls856 [Pdat-1::GFP, Pdat-1::LRRK2(G2019S), lin-15(+)]; wpdEx7 (pUnc-119::AGE1Orf::letuttr + pMyo-3::RFP) |  |
| DJM064 | age-1(hx546) II; cwrls856 [Pdat-1::GFP, Pdat-1::LRRK2(G2019S), lin-15(+)]; wpdEx8 (pUnc-119::AGE1Orf::letuttr + pMyo-3::RFP) |  |
| DJM065 | age-1(hx546) II; cwrls856 [Pdat-1::GFP, Pdat-1::LRRK2(G2019S), lin-15(+)]; wpdEx9 (pDat-1::AGE1Orf::let858 + pMyo-3::RFP) |  |
| DJM066 | age-1(hx546) II; cwrls856 [Pdat-1::GFP, Pdat-1::LRRK2(G2019S), lin-15(+)]; wpdEx10 (pDat-1::AGE1Orf::let858 + pMyo-3::RFP) |  |
| TU3401 | sid-1(pk3321) V; uls69[pCFJ90(myo-2p::mCherry) + unc-119p::sid-1] V | Calixto et al, 2010 |
| DJM001 | [Pdat-1::GFP, lin-15(+)]; sid-1(pk3321) V; uls69[pCFJ90(myo-2p::mCherry) + unc-119p::sid-1] V |  |
| DJM002 | [Pdat-1::GFP, Pdat-1::LRRK2(G2019S), lin-15(+)]; sid-1(pk3321) V; uls69[pCFJ90(myo-2p::mCherry) + unc-119p::sid-1] V |  |

## Acknowledgments

This work was supported by funding from the National Institutes of Health (R21AG058241) and Van Andel Institute. We thank Dr. Shu Chen (Case Western Reserve University) for providing the human LRRK2 transgenic strain. Some strains were provided by the CGC, which is funded by NIH Office of Research Infrastructure Programs (P40OD010440). We thank Dr. Jason Cooper (VAI) and Jennifer Kordich (VAI) for technical assistance in generating some worm strains.

## Supplemental figure legends

**Figure S1.**
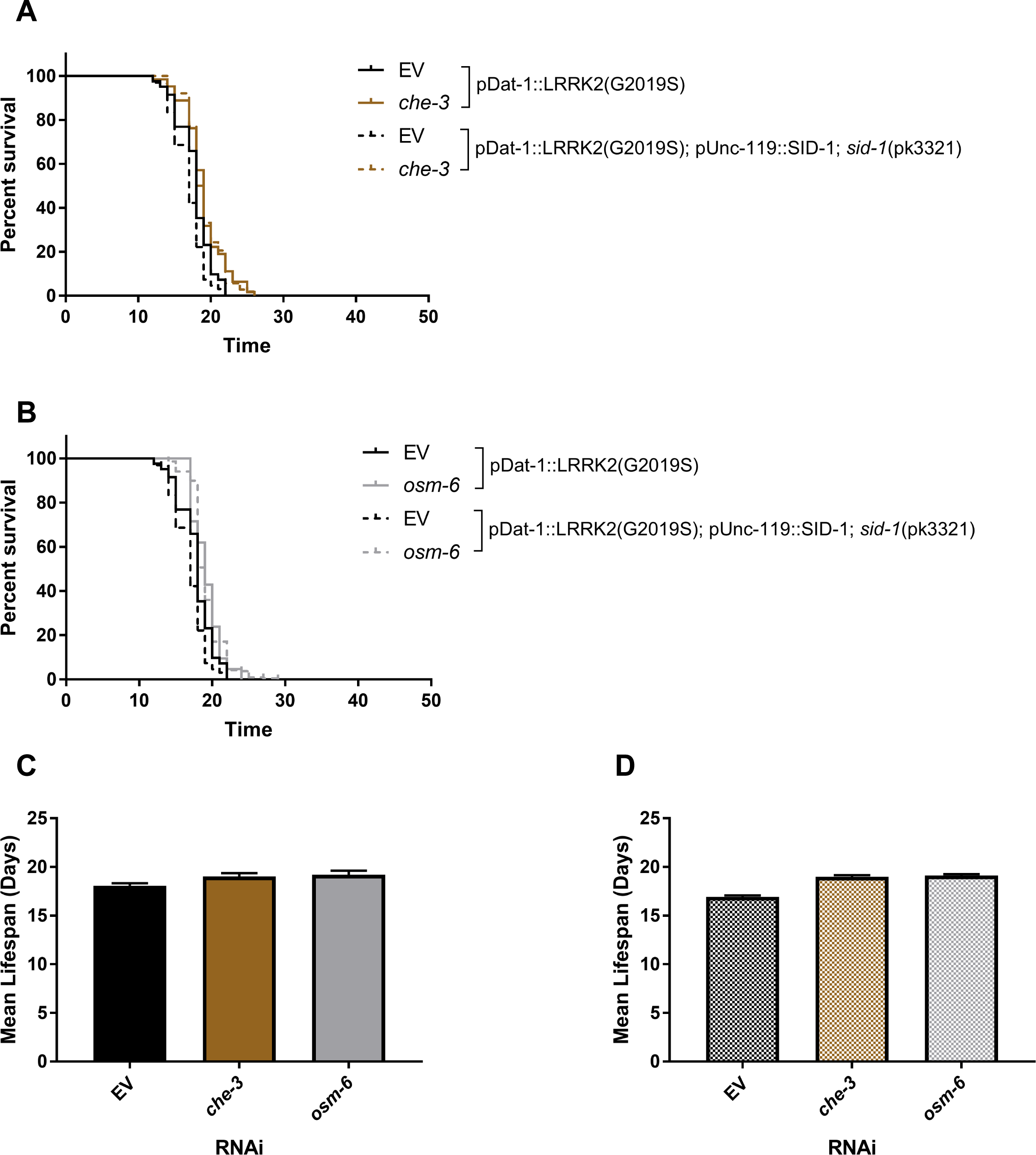
Systemic or neuronal-specific RNAi targeting chemosensory perception pathways fail to extend lifespan of LRRK2(G2019S) worms. An RNAi clone targeting *osm-5* was not available in the Ahringer RNAi library, therefore, we tested alternative candidates from this pathway for their ability to extend lifespan based on the work of Fujii et al, 2004. RNAi clones targeting *che-3* and *osm-6* both reveal extremely modest yet non-significant lifespan extension in systemic LRRK2(G2019S) or neuronal-specific LRRK2(G2019S); SID-1 worm strains at 100 µM FUdR. Shown are Kaplan-Meier survival curves (**A-B**) and mean lifespan for **C**) systemic or **D**) neuronal-specific *che-3* and *osm-5* RNAi. **C-D**) Bars represent mean ± SEM from ≥3 independent experiments (with *n* ≥ 30 animals/experiment). Differences between groups are not significant by one-way ANOVA with Dunnett’s post-hoc test.

**Figure S2.**
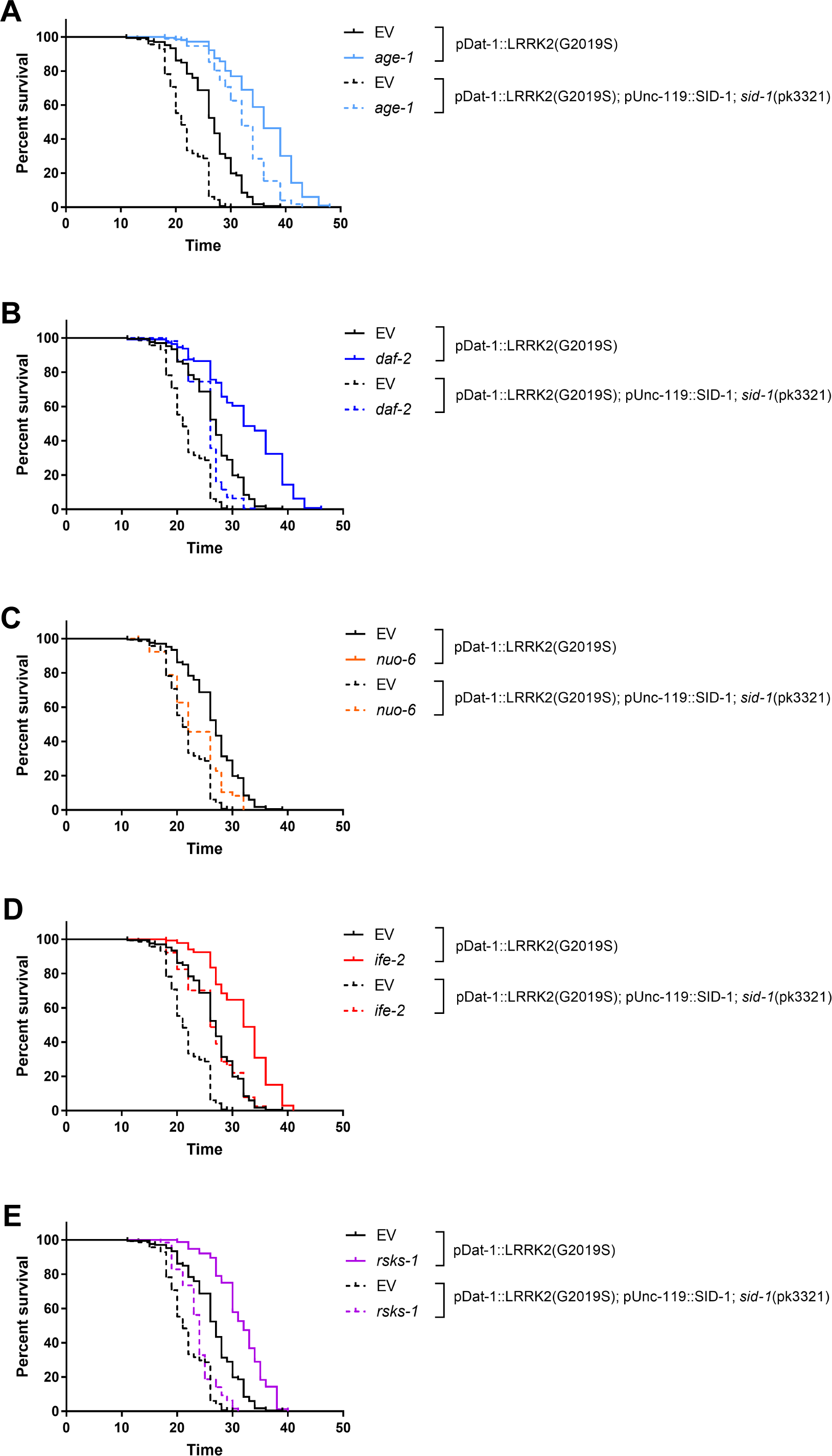
Kaplan-Meier survival curves comparing lifespan extension by systemic or neuronal-specific RNAi in LRRK2(G2019S) transgenic worms. Expression in the wild-type background confers systemic body-wide RNAi whereas a sensitized background [pUnc-119::SID-1; *sid-1(pk3321)*] confers neuronal-specific RNAi in LRRK2(G2019S) transgenic worms. Curves represent combined data from ≥3 individual experiments (*n* = 30 animals per experiment). Panneuronal or systemic knockdown of *age-1* and *ife-2* increase lifespan to similar extents, whereas neuronal-specific RNAi for *daf-2* and *rsks-1* extends lifespan to a lesser extent than systemic RNAi. Systemic RNAi for *nuo-6* is lethal.

**Figure S3.**
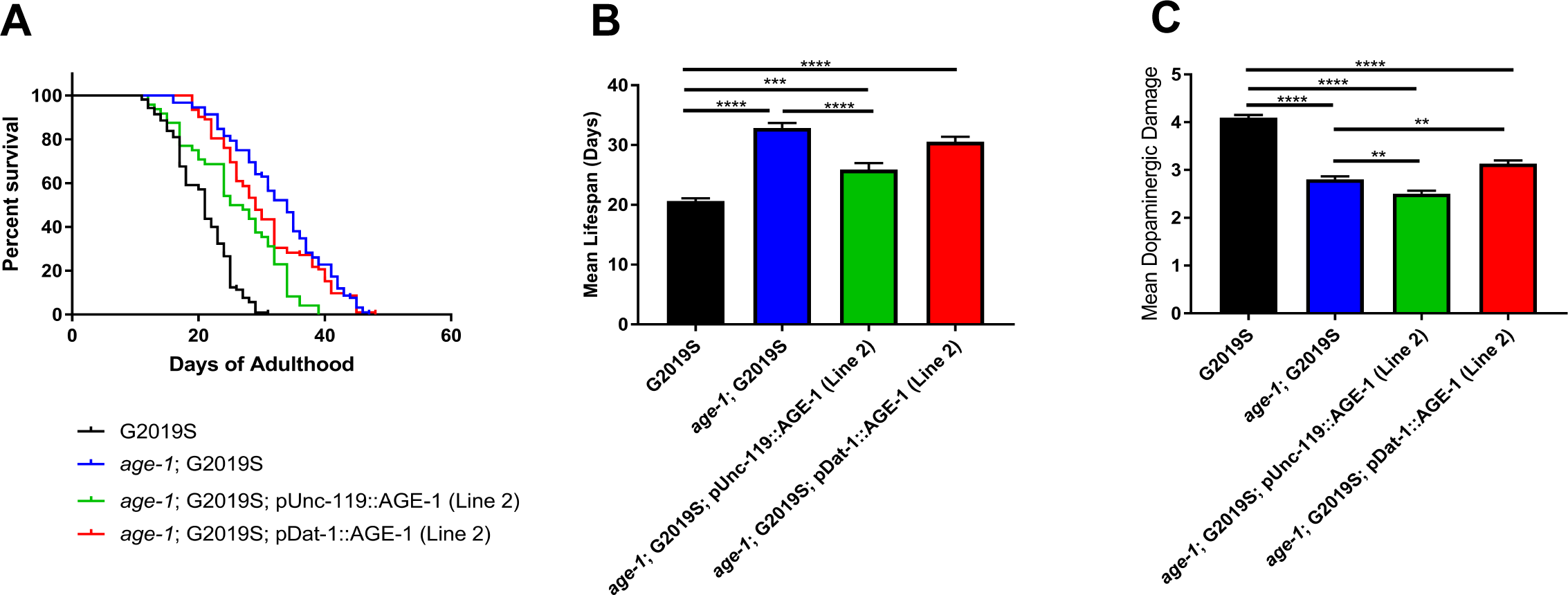
Lifespan and dopaminergic neuronal damage in LRRK2(G2019S); *age-1* worms with pan-neuronal or dopaminergic-specific expression of AGE-1/PI3K from independent transgenic arrays. **A**) Kaplan-Meier survival curves comparing the lifespan of LRRK2(G2019S) (black), *age-1*; LRRK2(G2019S) (blue), and independent transgenic arrays (line 2) with panneuronal (pUnc-119, green) or dopaminergic-specific (pDat-1, red) AGE-1 expression in *age-1*; LRRK2(G2019S) worms. Curves represent combined data from ≥3 individual experiments (*n* = 30 animals per experiment). **B**) Mean lifespan of each worm strain derived from (**A**). **C**) Quantification of dopaminergic neuronal damage at day 14 induced by LRRK2(G2019S) on a wild-type or *age-1(-)* background, or with tissue-specific restoration of AGE-1 on an *age-1(-)* background. **B-C**) Bars represent the mean ± SEM from ≥3 independent experiments with *n* ≥ 30 animals/experiment. \*\**P*<0.005, \*\*\**P*<0.001 or \*\*\*\**P*<0.0001 between indicated groups by oneway ANOVA with Tukey’s multiple comparisons post-hoc test.

